## Supplementary Tables and Figures for "The *Listeria monocytogenes* persistence factor ClpL is a potent stand-alone disaggregase"

Table S1: strains and plasmids used in this study

| Strain | Description | Source or reference |
|---|---|---|
| *E. coli* XL1 blue | *recA1 endA1 gyrA96 thi-1 hsdR1 supE44 relA1 lac* [F' *proAB* $lacI^q$ *ΔM15* Tn10 (Tcr)] | Stratagene |
| *E. coli* BL21 | *F- ompT lon hsdSB gal dcm* $\lambda$ (DE3) | Novagen |
| *E. coli ΔclpB* | MC1000 Δ*clpB*::Km | (1) |
| *E. coli dnak103* | MC4100 *dnak103* | (1) |
| **Plasmid** | **Description** | **Source or reference** |
| pET24a-*clpL* | Vector for IPTG-inducible expression of *clpL* in *E.coli* BL21 cells, fused C-terminal 6xHis-tag | This study |
| pET24a-*clpL-ΔN* | Vector for IPTG-inducible expression of *clpL-ΔN* in *E.coli* BL21 cells, fused C-terminal 6xHis-tag | This study |
| pET24a-*Lm_clpB* | Vector for IPTG-inducible expression of *Lm clpB* in *E.coli* BL21 cells, fused C-terminal 6xHis-tag | This study |
| pET24a-*Lm_DnaK* | Vector for IPTG-inducible expression of *Lm_DnaK* in *E.coli* BL21 cells, fused C-terminal 6xHis-tag | This study |
| pET24a-*Lm_DnaJ* | Vector for IPTG-inducible expression of *Lm_DnaJ* in *E.coli* BL21 cells, fused C-terminal 6xHis-tag | This study |
| pET24a-*Lm_GrpE* | Vector for IPTG-inducible expression of *Lm_GrpE* in *E.coli* BL21 cells, fused C-terminal 6xHis-tag | This study |
| pET24a-*clpL-Patch_A* | Vector for IPTG-inducible expression of *clpL-Patch_A* in *E.coli* BL21 cells, fused C-terminal 6xHis-tag | This study |
| pET24a-*clpL-Patch_B* | Vector for IPTG-inducible expression of *clpL-Patch_B* in *E.coli* BL21 cells, fused C-terminal 6xHis-tag | This study |
| pET24a-*clpL-Patch_C* | Vector for IPTG-inducible expression of *clpL-Patch_C* in *E.coli* BL21 cells, fused C-terminal 6xHis-tag | This study |
| pET24a-*clpL-Patch_D* | Vector for IPTG-inducible expression of *clpL-Patch_D* in *E.coli* BL21 cells, fused C-terminal 6xHis-tag | This study |
| pET24a-*clpL-Patch_E* | Vector for IPTG-inducible expression of *clpL-Patch_E* in *E.coli* BL21 cells, fused C-terminal 6xHis-tag | This study |
| pET24a-*clpL-Patch_AB* | Vector for IPTG-inducible expression of *clpL-Patch_AB* in *E.coli* BL21 cells, fused C-terminal 6xHis-tag | This study |
| pET24a-*clpL-Patch_AC* | Vector for IPTG-inducible expression of *clpL-Patch_AC* in *E.coli* BL21 cells, fused C-terminal 6xHis-tag | This study |
| pET24a-*clpL-Patch_BC* | Vector for IPTG-inducible expression of *clpL-Patch_BC* in *E.coli* BL21 cells, fused C-terminal 6xHis-tag | This study |
| pET24a-*clpL_all_NQ→Ala* | Vector for IPTG-inducible expression of *clpL-Patch_all_NQ→Ala* in *E.coli* BL21 cells, fused C-terminal 6xHis-tag | This study |
| pET24a-*clpL-Patch_all_Aro→Ala* | Vector for IPTG-inducible expression of *clpL-Patch_all_Aro→Ala* in *E.coli* BL21 cells, fused C-terminal 6xHis-tag | This study |
| pET24a-*clpL-Y36A* | Vector for IPTG-inducible expression of *clpL-Y36A* in *E.coli* BL21 cells, fused C-terminal 6xHis-tag | This study |
| pET24a-*clpL-F48A* | Vector for IPTG-inducible expression of *clpL-F48A* in *E.coli* BL21 cells, fused C-terminal 6xHis-tag | This study |
| pET24a-*clpL-Y51A* | Vector for IPTG-inducible expression of *clpL-Y51A* in *E.coli* BL21 cells, fused C-terminal 6xHis-tag | This study |
| pET24a-*clpL-NTD61* | Vector for IPTG-inducible expression of *clpL_NTD61* in *E.coli* BL21 cells, fused C-terminal 6xHis-tag | This study |
| pET24a-*clpL-NTD78* | Vector for IPTG-inducible expression of *clpL_NTD78* in *E.coli* BL21 cells, fused C-terminal 6xHis-tag | This study |
| pCA528-*clpL-E352A* | Vector for IPTG-inducible expression of *clpL-E352A* in *E.coli* BL21 cells, fused cleavable N-terminal 6xHis-SUMO-tag | This study |
| pCA528-*clpL-F354A* | Vector for IPTG-inducible expression of *clpL-F354A* in *E.coli* BL21 cells, fused cleavable N-terminal 6xHis-SUMO-tag | This study |
| pCA528-*clpL-T355C* | Vector for IPTG-inducible expression of *clpL-T355C* in *E.coli* BL21 cells, fused cleavable N-terminal 6xHis-SUMO-tag | This study |
| pC6AmpΔBsaI-$L_N$-*ClpB** | Vector for IPTG-inducible expression of $L_N$-*ClpB** in *E.coli* BL21 cells, fused cleavable N-terminal 6xHis-SUMO-tag | This study |
| pC6AmpΔBsaI-$L_N$-*ClpB*-E199A-E598A* | Vector for IPTG-inducible expression of $L_N$-*ClpB*- E199A-E598A* in *E.coli* BL21 cells, fused cleavable N-terminal 6xHis-SUMO-tag | This study |
| pC6AmpΔBsaI-$L_N$-*ClpB*-Patch_A* | Vector for IPTG-inducible expression of $L_N$-*ClpB*-Patch_A* in *E.coli* BL21 cells, fused cleavable N-terminal 6xHis-SUMO-tag | This study |
| pC6AmpΔBsaI-$L_N$-*ClpB*-Patch_AB* | Vector for IPTG-inducible expression of $L_N$-*ClpB*-Patch_AB* in *E.coli* BL21 cells, fused cleavable N-terminal 6xHis-SUMO-tag | This study |
| pC6AmpΔBsaI-$L_N$-*ClpB*-Patch_C* | Vector for IPTG-inducible expression of $L_N$-*ClpB*-Patch_C* in *E.coli* BL21 cells, fused cleavable N-terminal 6xHis-SUMO-tag | This study |

| | | |
|---|---|---|
| pC6AmpΔBsaI-$L_N$-ClpB*-Y36A | Vector for IPTG-inducible expression of $L_N$-ClpB*-Y36A in *E.coli* BL21 cells, fused cleavable N-terminal 6xHis-SUMO-tag | This study |
| pC6AmpΔBsaI-$L_N$-ClpB*-F48A | Vector for IPTG-inducible expression of $L_N$-ClpB*-F48A in *E.coli* BL21 cells, fused cleavable N-terminal 6xHis-SUMO-tag | This study |
| pC6AmpΔBsaI-$L_N$-ClpB*-Y51A | Vector for IPTG-inducible expression of $L_N$-ClpB*-Y51A in *E.coli* BL21 cells, fused cleavable N-terminal 6xHis-SUMO-tag | This study |
| pUHE21-2fd12 | Empty Vector control for *in vivo* assays | This study |
| pUHE21-$clpG_{GI}$ | Vector for IPTG-inducible expression of $clpG_{GI}$ in *E.coli ΔclpB* cells | adjust |
| pUHE21-*Ec clpB* | Vector for IPTG-inducible expression of *Ec clpB* in *E.coli ΔclpB* cells | adjust |
| pDS56-*clpL* | Vector for IPTG-inducible expression of *clpL* in *E.coli* □*clpB* cells and *E. coli dnak103* cells | This study |
| pUHE21-*clpL-ΔN* | Vector for IPTG-inducible expression of *clpL* in *E.coli* □*clpB* cells and *E. coli dnak103* cells | This study |
| pDS56-*clpL-E352A* | Vector for IPTG-inducible expression of *clpL* in *E.coli* □*clpB* cells | This study |
| pDS56-*clpL-F354A* | Vector for IPTG-inducible expression of *clpL* in *E.coli* □*clpB* cells | This study |
| pUHE21-*Ec* dnaK | Vector for IPTG-inducible expression of *Ec dnaK* in *E.coli dnak103* cells | (1) |
| pET24a-$clpG_{GI}$ | Vector for IPTG-inducible expression of $clpG_{GI}$ in *E.coli* BL21 cells, fused C-terminal 6xHis-tag | adjust |
| pDS56-*Ec clpB* | Vector for IPTG-inducible expression of *Ec clpB* in *E.coli ΔclpB* cells, fused C-terminal 6xHis-tag | (1) |
| pUHE21-*Ec dnaK*-His | Vector for IPTG-inducible expression of *Ec dnaK* in *E.coli ΔclpB* cells, fused C-terminal 6xHis-tag | (1) |
| pCA528-*Ec dnaJ* | Vector for IPTG-inducible expression of *Ec dnaJ* in *E.coli* BL21 cells, fused cleavable N-terminal 6xHis-SUMO-tag | (1) |
| pCA528-*Ec grpE* | Vector for IPTG-inducible expression of *Ec grpE* in *E.coli* BL21 cells, fused cleavable N-terminal 6xHis-SUMO-tag | (1) |
| pDS56-*Ec* ΔN-ClpB-K476C | Vector for IPTG-inducible expression of *Ec ΔN-clpB-K476C* in *E.coli ΔclpB* cells, fused C-terminal 6xHis-tag | (2) |
| pDS56-*luciferase* | Vector for IPTG-inducible expression of *luciferase* in *E.coli ΔclpB* cells, fused N-terminal 6xHis-tag | (3) |
| pDS56-*luciferase-yfp* | Vector for IPTG-inducible expression of *luciferase-YFP* in *E.coli ΔclpB* cells, fused N-terminal 6xHis-tag | (3) |

| **Table S2: number of replicates** | | | | | | | | | |
|---|---|---|---|---|---|---|---|---|---|
| # | 1 | 2 | 3 | 4 | 5 | 6 | 7 | 8 | 9 |
| number of replicates by figure by sample in plot order (left-right or top-down, according to labels) | | | | | | | | | |
| Fig. 1b1 | 58 | | | | | | | | |
| Fig. 1b2 | 18 | 18 | 8 | 4 | 10 | 4 | 4 | | |
| Fig. 1c | 3 | 4 | 3 | 3 | 3 | | | | |
| Fig. 1e | 6 | 5 | | | | | | | |
| Fig. 2b | 3 | 3 | 4 | 4 | 3 | | | | |
| Fig. 2c | 4 | 4 | 4 | 4 | | | | | |
| Fig. 2d | 3 | 3 | 3 | | | | | | |
| Fig. 3a | 8 | 8 | 5 | 5 | | | | | |
| Fig. 3b | 3 | 3 | | | | | | | |
| Fig. 3c | 4 | 4 | 4 | 4 | 4 | 4 | 4 | | |
| Fig. 3e | 3 | 3 | 3 | 3 | 3 | 3 | 3 | | |
| Fig. 3f | 4 | 4 | 4 | 4 | 4 | 4 | 4 | | |
| Fig. 4e | 29 | 28 | 4 | 6 | 13 | 3 | 3 | 5 | 20 |
| Fig. 4f | 9 | 9 | 4 | 4 | 7 | 3 | 3 | 3 | 7 |
| Fig. 4g | 3 | 3 | 3 | 3 | | | | | |
| Fig. 5b | 13 | 13 | 13 | 13 | 13 | 13 | 13 | | |
| Fig. 5c | 3 | 3 | 3 | 3 | | | | | |
| Fig. 5d | 3 | 3 | 3 | 3 | | | | | |
| Fig. 6c (particles, not replicates) | 5233 | 18314 | 1900 | 14215 | 7074 | 11765 | | | |
| Fig. 6d | 10 | 6 | 4 | 4 | | | | | |
| Fig. 6e | 8 | 5 | 3 | 3 | | | | | |
| Fig. 6g | 6 | 6 | 6 | 6 | 6 | 6 | | | |
| Fig. 1 – figure s2a | 3 | 3 | 3 | 3 | | | | | |
| Fig. 1 – figure s2c | 4 | 3 | 4 | 3 | | | | | |
| Fig. 1 – figure s2d | 5 | 5 | 5 | 5 | 5 | | | | |
| Fig. 1 – figure s2e (left) | 19 | | | | | | | | |
| Fig. 1 – figure s2e (right) | 3 | 3 | 3 | 3 | 3 | 3 | 3 | | |
| Fig. 1 – figure s2f | 13 | 6 | 4 | 4 | 3 | 3 | 6 | n.d. | |
| Fig. 1 – figure s2g | 5 | 5 | 5 | | | | | | |
| Fig. 2 – figure s1 | 5 | 4 | 4 | 3 | 4 | 4 | 3 | | |

| | | | | | | | | |
|---|---|---|---|---|---|---|---|---|
| Fig. 3 – figure s1b | 24 | 19 | | | | | | |
| Fig. 3 – figure s1c | 3 | 3 | 3 | 3 | 3 | 3 | | |
| Fig. 3 – figure s1d | 3 | 3 | 3 | 3 | 3 | 3 | 3 | |
| Fig. 3 – figure s2d | 3 | 3 | | | | | | |
| Fig. 4 – figure s2a (left) | 12 | 9 | 3 | 12 | 3 | 3 | 3 | 7 |
| Fig. 4 – figure s2a (right) | 3 | 3 | 3 | 3 | | | | |
| Fig. 4 – figure s2b | 24 | 22 | 7 | 7 | 9 | 9 | 9 | |
| Fig. 4 – figure s2c | 10 | 8 | 5 | 5 | 5 | 5 | 5 | |
| Fig. 4 – figure s2d (left) | 9 | 9 | 6 | 6 | 3 | 3 | 6 | |
| Fig. 4 – figure s2d (right) | 7 | 3 | 4 | 4 | 4 | | | |
| Fig. 4 – figure s2e | 14 | 14 | 10 | 10 | 10 | 6 | 6 | 6 |
| Fig. 4 – figure s2f | 3 | 3 | 3 | 3 | 3 | 3 | 3 | 3 |
| Fig. 4 – figure s2g (left) | 3 | 3 | 3 | 3 | 3 | | | |
| Fig. 4 – figure s2g (right) | 7 | 7 | 3 | 3 | 3 | 4 | 4 | 4 |
| Fig. 4 – figure s2h | 3 | 3 | 3 | 3 | 5 | 3 | | |
| Fig.5 - figure s1a | 3 | 3 | 3 | 3 | 3 | 3 | | |
| Fig.5 - figure s1b | 3 | 3 | 3 | 3 | | | | |
| Fig.5 - figure s1c (black) | 13 | 13 | 13 | 13 | 13 | 13 | 13 | |
| Fig.5 - figure s1c (red) | 13 | 13 | 13 | 13 | 13 | 13 | 13 | |
| Fig. 5 – figure s1d (red) | 13 | 13 | 13 | 13 | 13 | 13 | 13 | |
| Fig. 5 – figure s1e (black) | 13 | 13 | 13 | 13 | 13 | 13 | 13 | |
| Fig. 5 – figure s1e (red) | 13 | 13 | 13 | 13 | 13 | 13 | 13 | |
| Fig. 6 – figure s1d | 3 | 3 | 3 | 4 | 4 | 4 | | |
| Fig. 6 – figure s1e (particles, not replicates) | 3588 | 7404 | 9655 | | | | | |
| Fig. 6 – figure s1f (left) | 6 | 3 | 3 | 3 | | | | |
| Fig. 6 – figure s1f (right) | 6 | 3 | 3 | 3 | | | | |
| Fig. 6 – figure s2a (left) | 3 | 3 | | | | | | |
| Fig. 6 – figure s2a (right) | 3 | 3 | | | | | | |
| Fig. 6 – figure s2b | 7 | 7 | | | | | | |
| Fig. 6 – figure s2d | 6 | 6 | 6 | 6 | 6 | 6 | | |

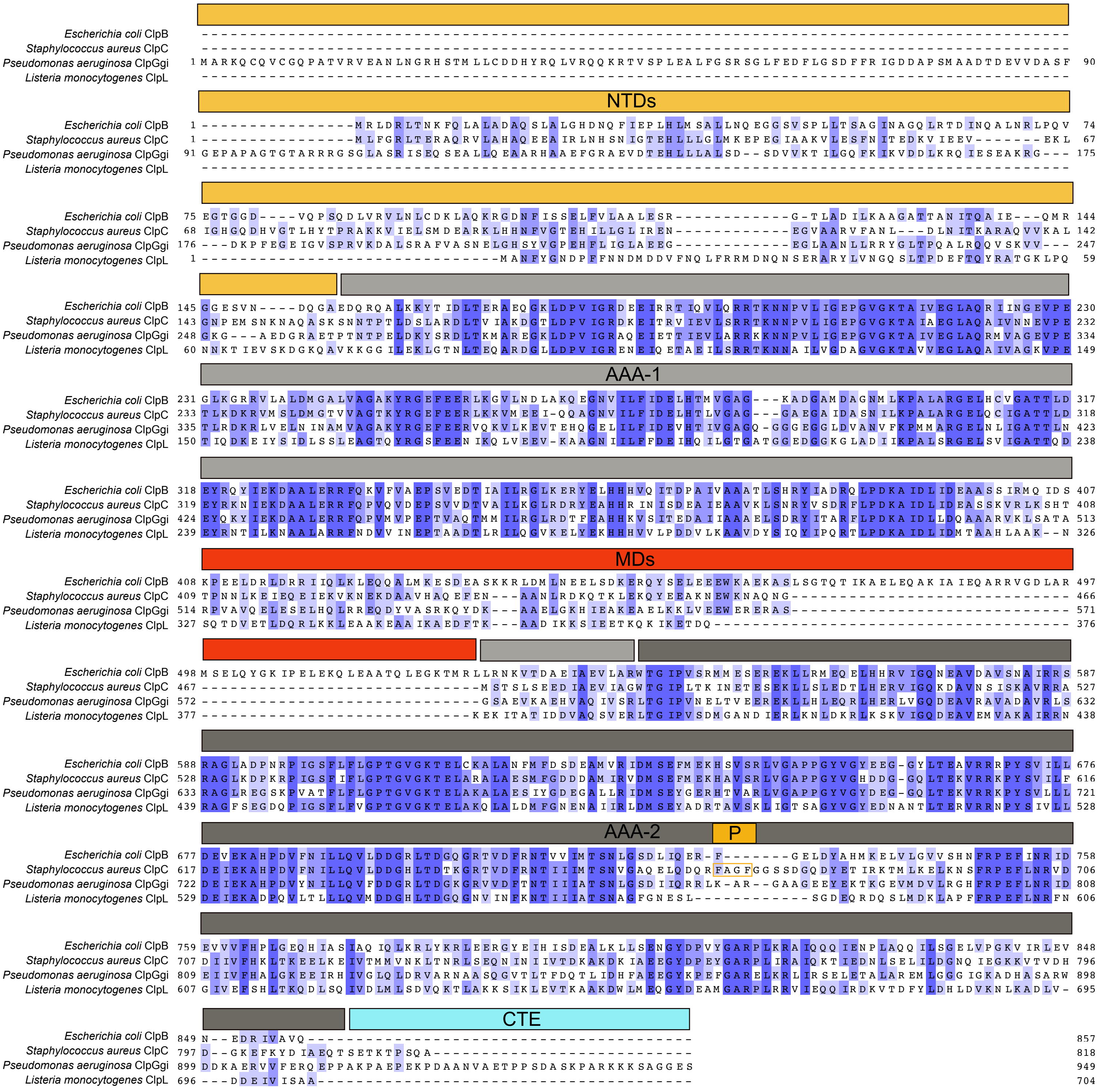

**Figure 1 - figure supplement 1**

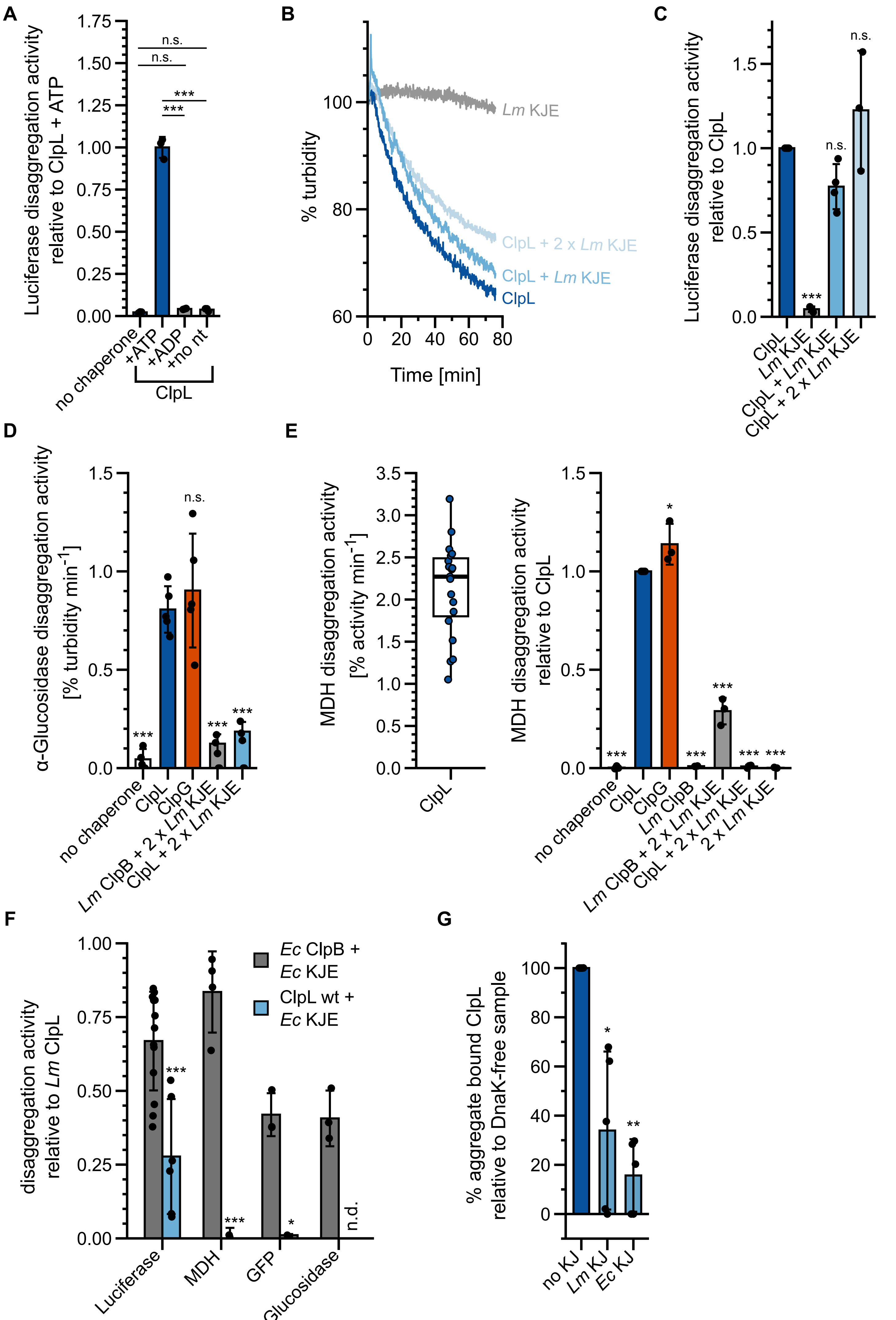

Figure 1 - figure supplement 2

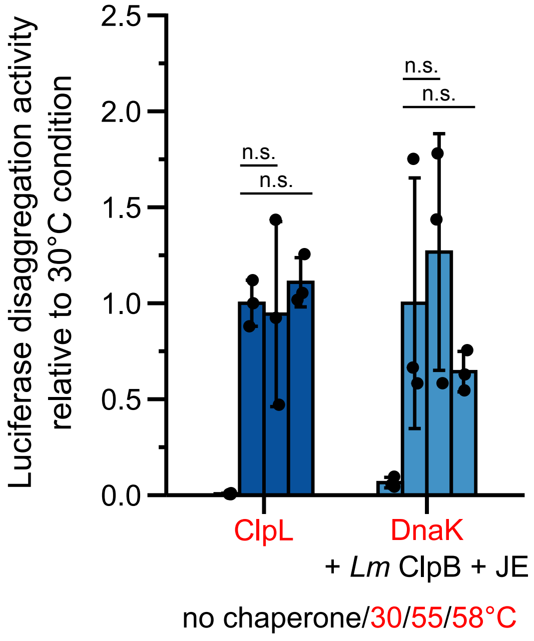

**Figure 2 - figure supplement 1**

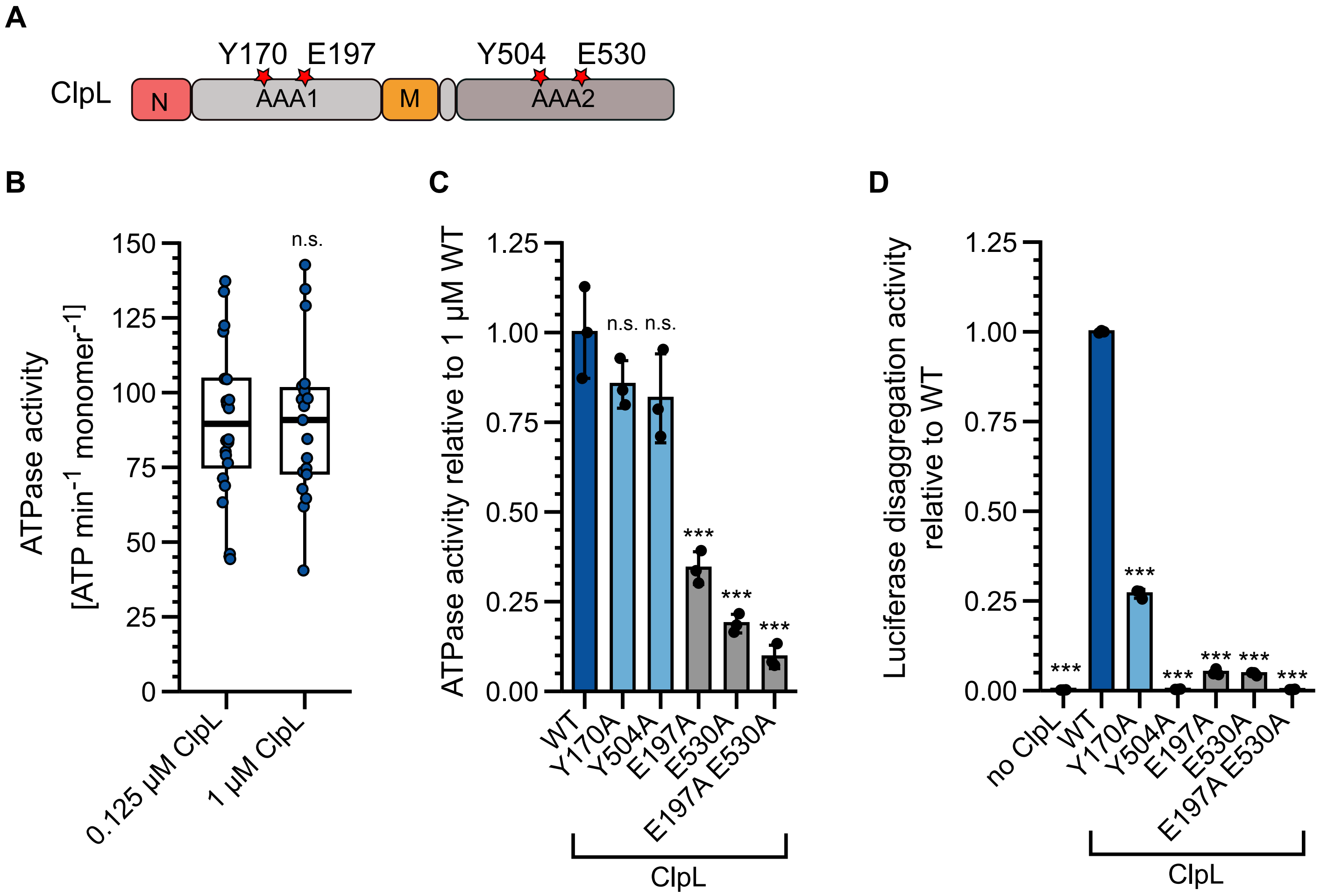

**Figure 3 - figure supplement 1**

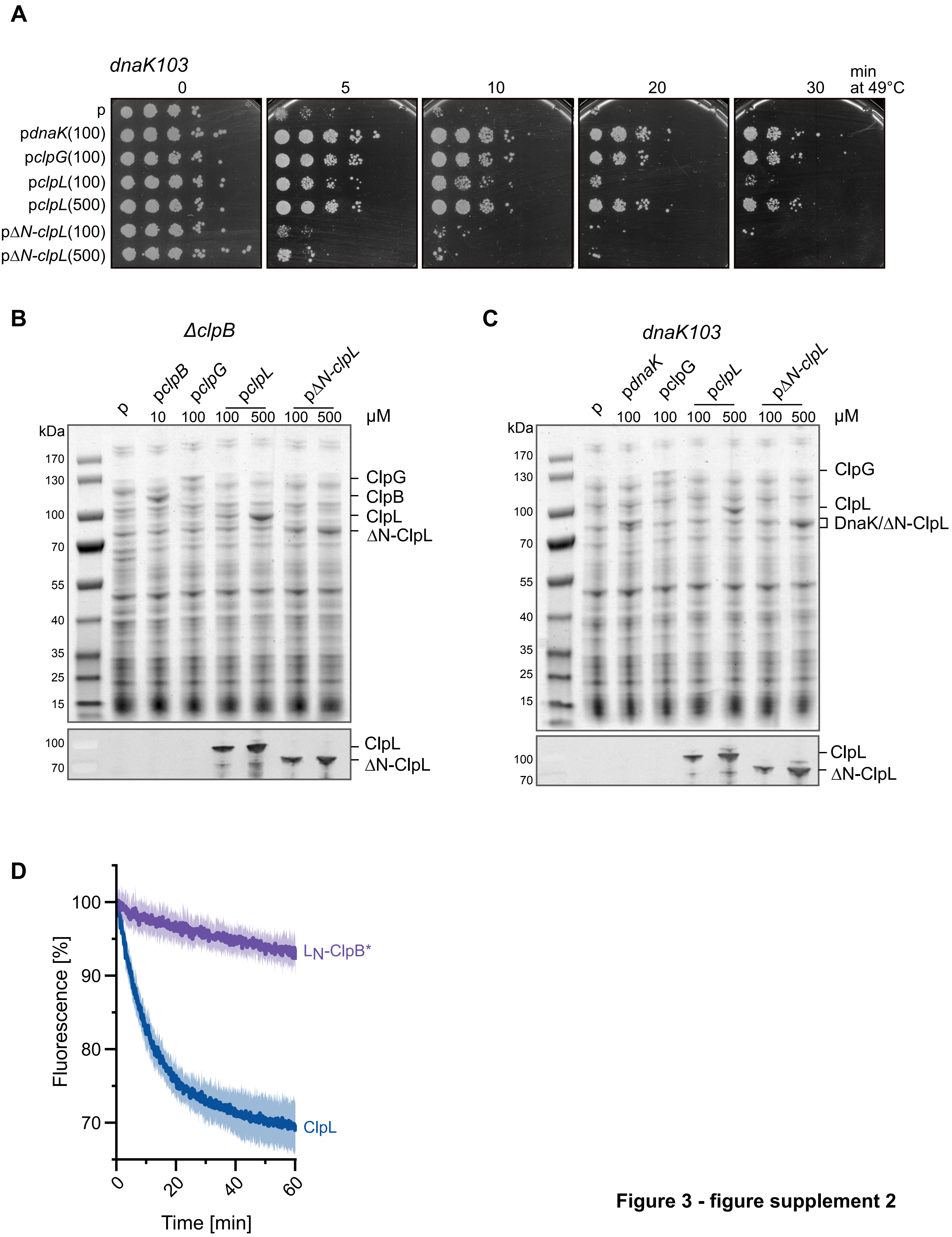

**Figure 3 - figure supplement 2**

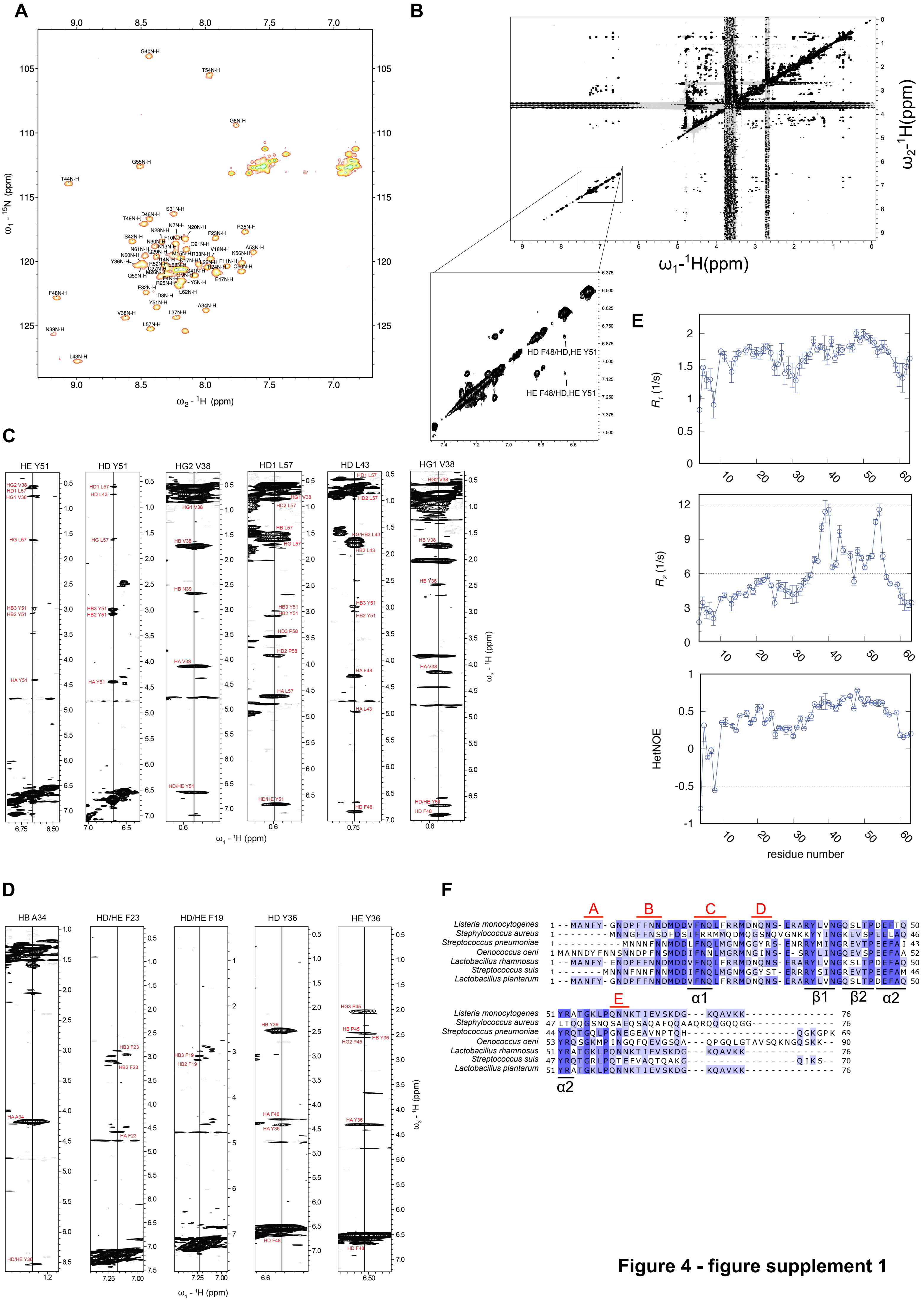

Figure 4 - figure supplement 1

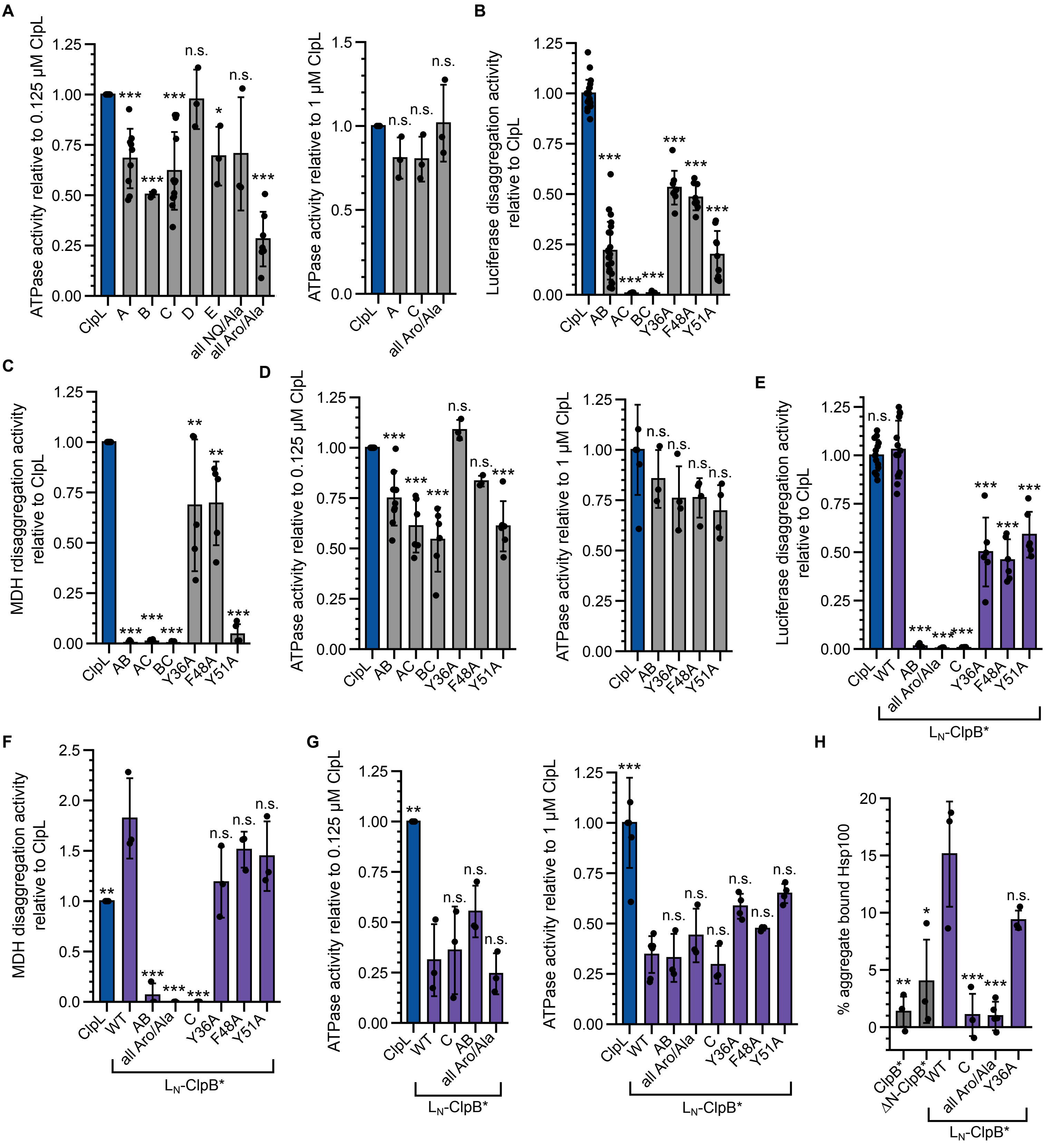

**Figure 4 - figure supplement 2**

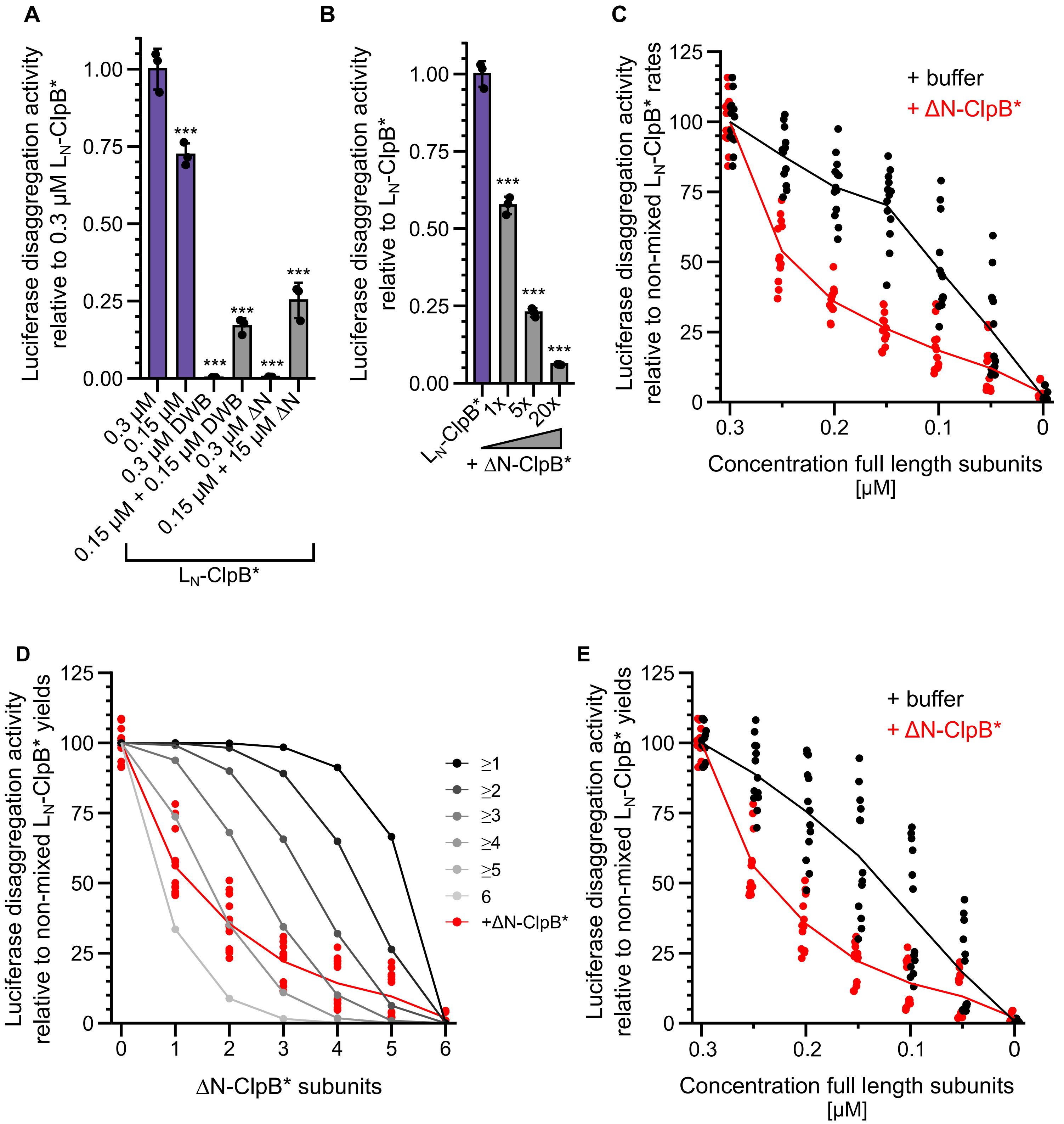

**Figure 5 - figure supplement 1**

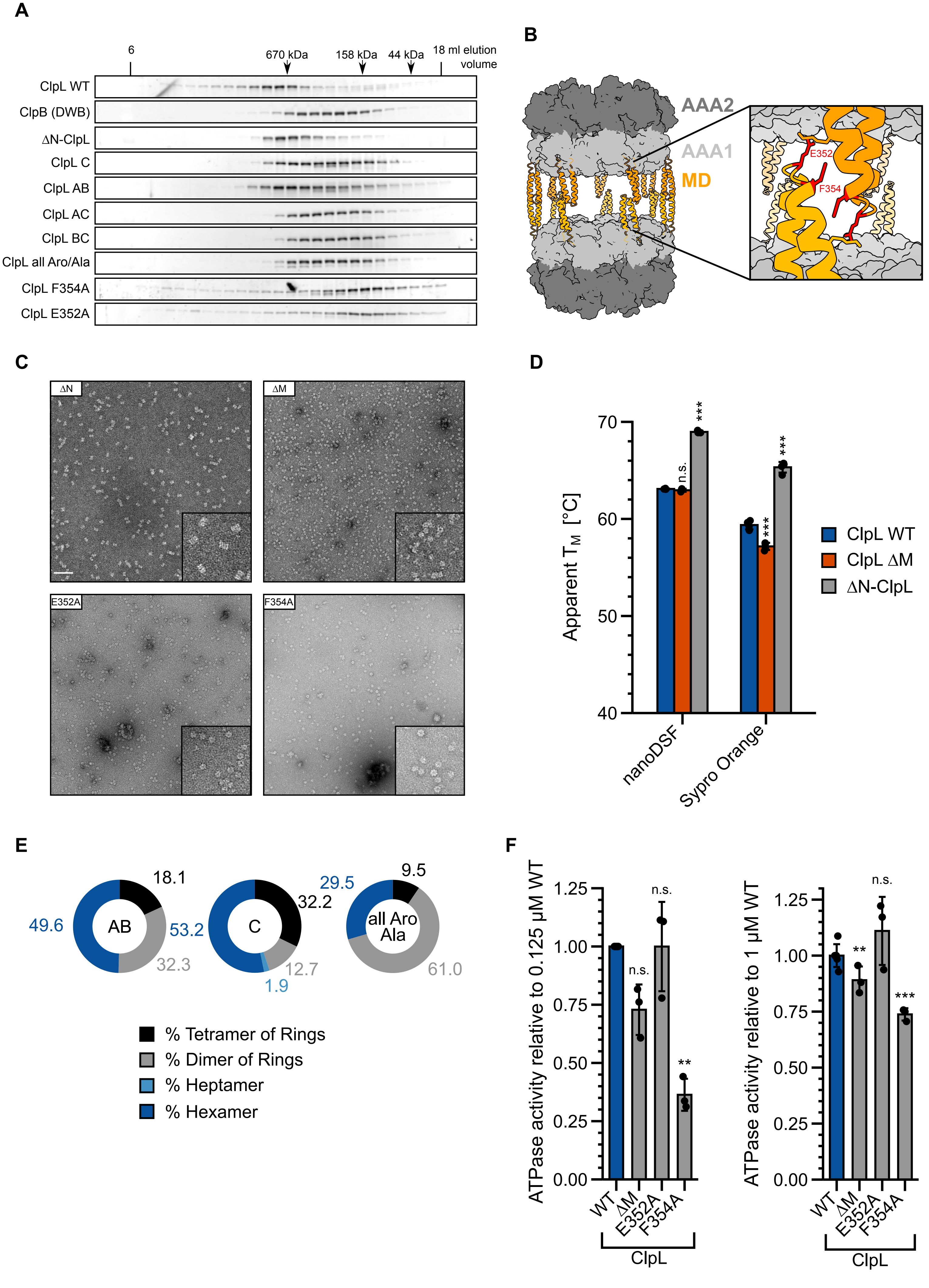

Figure 6 - figure supplement 1

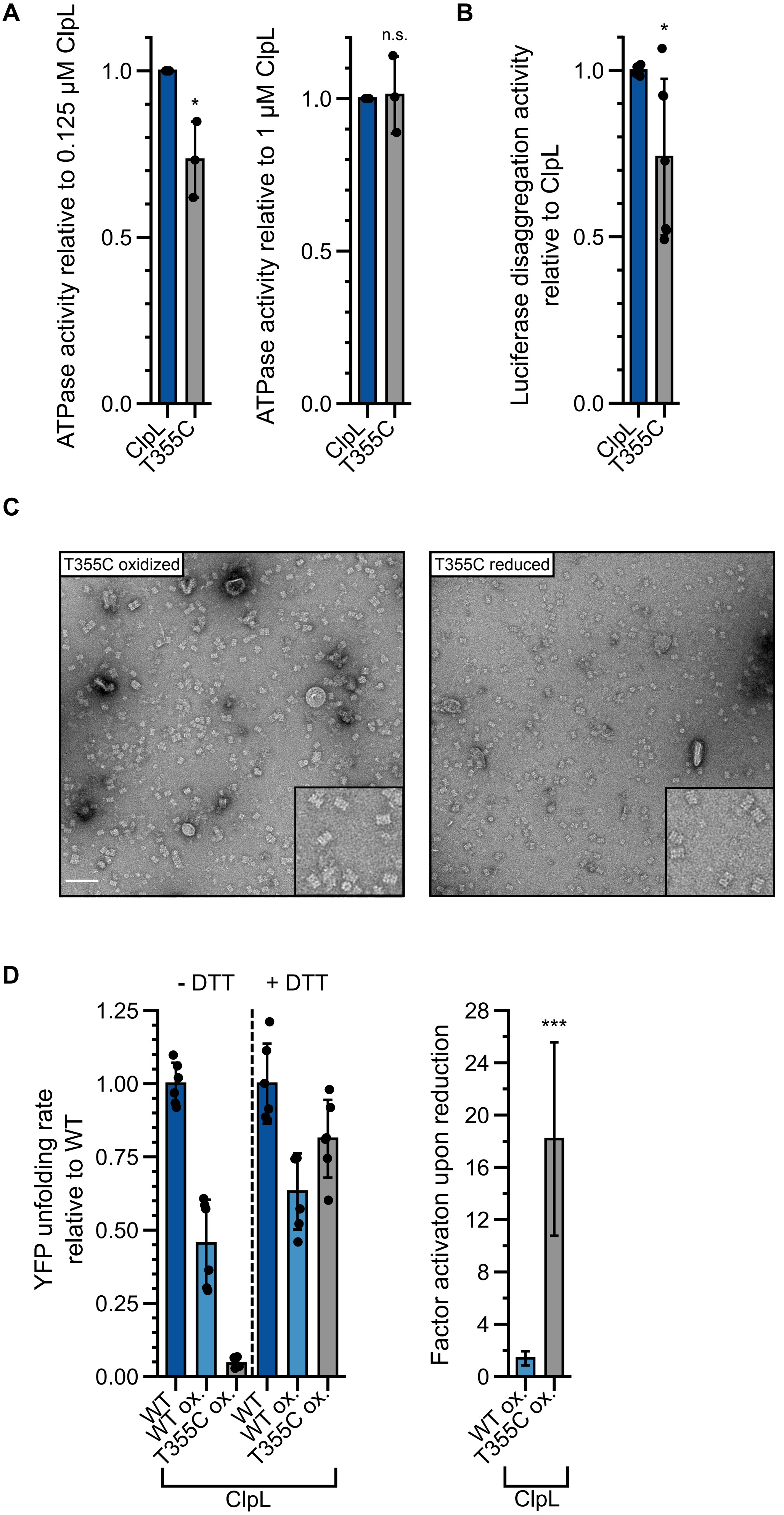

**Figure 6 - figure supplement 2**

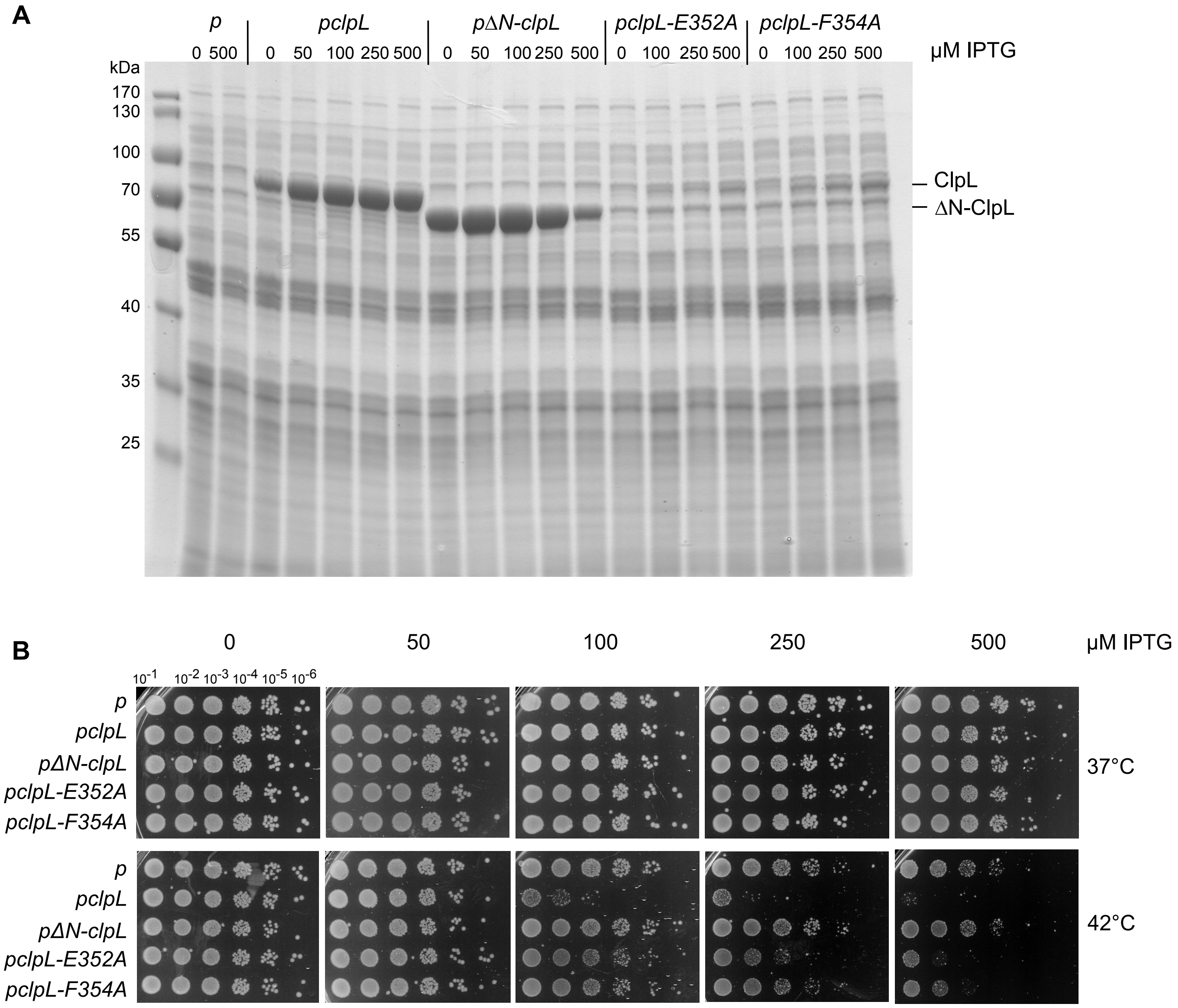

**Figure 6 - figure supplement 3**

**Supplementary figure legends**

**Figure 1 – figure supplement 1**
Sequence alignment of *Escherichia coli* ClpB, *Staphylococcus aureus* ClpC, *Pseudomonas aeruginosa* $ClpG_{GI}$ and *Listeria monocytogenes* ClpL. The domain organization is indicated. Similar and identical residues are highlighted in light and dark blue.

**Figure 1 – figure supplement 2**
ClpL is a potent, stand-alone disaggregase. (A) Relative Luciferase disaggregation activities (% refolded Luciferase/min) of ClpL in presence of 2 mM ATP (and a regeneration-mix), 2 mM ADP and without nucleotide were determined. The disaggregation activity of ClpL WT (+ ATP) was set to 1. (B) Disaggregation of aggregated Luciferase was monitored by turbidity measurements in presence of indicated chaperones. KJE: DnaK/DnaJ/GrpE. Initial turbidity was set to 100%. (C) Relative Luciferase disaggregation activities (% turbidity/min) were determined for indicated chaperone combinations. The disaggregation activity of ClpL was set to 1. (D) Relative α-Glucosidase disaggregation activities (% turbidity/min) were determined in presence of indicated chaperones. Initial turbidity of α-Glucosidase aggregates was set to 100%. Disaggregation activity of ClpL was set to 1. (E) Left: MDH disaggregation activity of ClpL (% refolded MDH/min). Right: Relative MDH disaggregation activities of indicated chaperones were determined. The activity of ClpL was set to 1. (F) Relative disaggregation activities of indicated chaperone combinations towards aggregated Luciferase, MDH, GFP and α-Glucosidase were determined. The activity of ClpL was set to 1. n.d.: not determined. (G) ATPase-deficient ClpL-E197A/E530A) with and without the *Lm/Ec* DnaK system (KJ) was incubated with aggregated MDH in presence of ATP. The extent of aggregate binding (% total protein) was determined by co-sedimentation upon centrifugation. Percentage of aggregate bound ClpL without KJ was set to 1. Shown are representative curves (A), a boxplot (D) or data points and mean ± SD (B/C/D/E), n ≥ 3. Statistical Analysis: One-way ANOVA, Welch's Test for post-hoc multiple comparisons. Significance levels: *p < 0.05; **p < 0.01; ***p < 0.001. n.s.: not significant.

**Figure 2 – figure supplement 1**
Unfolding of DnaK at high temperatures is reversible. Relative Luciferase disaggregation activities (% refolded Luciferase/min) of ClpL and *Lm* ClpB/KJE were determined. ClpL or DnaK were incubated for 30 min at the indicated temperatures prior to the Luciferase disaggregation performed at 30°C. The 30°C condition was set to 1 for each chaperone respectively.Shown are mean curves ± SD (A) or data points and mean ± SD (B), n ≥ 3. Statistical Analysis: One-way ANOVA, Welch's Test for post-hoc multiple comparisons. Significance levels: *p < 0.05; **p < 0.01; ***p < 0.001. n.s.: not significant.

**Figure 3 – figure supplement 1**
ClpL disaggregation activity relies on ATP-fueled substrate threading. (A) Domain organization of ClpL. Pore loop (Y170, Y504) and Walker B (E197, E530) residues of AAA1 and AAA2 domains are indicated. (B) ClpL ATPase activities were determined at 0.125 and 1 µM protein concentrations. (C) Relative ATPase activities of ClpL WT and indicated mutants were determined at 1 µM protein concentration. The ATPase activity of ClpL WT was set to 1. (D) Relative Luciferase disaggregation activities (% refolded Luciferase/min) of ClpL WT and indicated mutants were determined. The activity of ClpL WT was set to 1. Shown are boxplots (B) or data points and mean ± SD (C/D), n ≥ 3. Statistical Analysis: One-way ANOVA, Welch's Test for post-hoc multiple comparisons. Significance levels: *p < 0.05; **p < 0.01; ***p < 0.001. n.s.: not significant.

**Figure 3 – figure supplement 2**

The ClpL NTD is crucial for disaggregation activity. (A) *E. coli dnaK103* cells harboring plasmids for IPTG-controlled expression of indicated disaggregases were grown at 30°C to mid-logarithmic growth phase and shifted to 49°C. Serial dilutions of cells were prepared at indicated time points, spotted on LB plates and incubated at 30°C. 100/500: μM IPTG added to induce disaggregase expression. p: empty vector control. (B-C) Production levels of disaggregases (ClpB, DnaK, ClpG, ClpL, ClpL-ΔN) in *E. coli ΔclpB* (B) and *dnaK103* (C) mutant cells. Cells were were grown at 30°C for 1.5 h and disaggregase expression was induced by addition of indicated IPTG concentrations (μM) for 2 h. Total cell extracts were prepared and levels of disaggregases were determined SDS-PAGE followed by Coomassie-staining. ClpL levels were additionally determined by western blot analysis using ClpL-specific antibodies. *p*: empty vector control. (D) The ClpL NTD does not increase ClpB* threading power to ClpL levels. Aggregated Luciferase-YFP was incubated in presence of ClpL or $L_N$-ClpB* and YFP fluorescence was recorded. Initial YFP fluorescence was set at 100%. Shown are mean curves ± SD, $n \geq 3$.

**Figure 4 – figure supplement 1**

Structural analysis of ClpL NTD. (A) $^1$H,$^{15}$N-HSQC of ClpL NTD with resonance assignment labeled at each peak. Several residues exhibit dispersed peak position, indicative of folding. (B) $^1$H,$^1$H-2D NOESY in $D_2O$ of ClpL NTD confirming NOEs between aromatic ring protons of the hydrophobic core formed by α2 and the β-sheet. NOEs of the second predicted core between α1 and α2 are absent, indicating that there is no interaction between α1 and α2. (C) Selected strips of the $^{13}$C-edited 3D NOESY-HSQC to illustrate that the predicted hydrophobic core between α2 and the β-sheet is present. Multiple cross-peaks (NOEs) are present between relevant residues (Y36, V38, L43, F48, Y51). (D) Selected strips of the $^{13}$C-edited 3D NOESY-HSQC illustrating the absence of NOEs expected to between residues of the second hydrophobic core between α1 and α2. For these residues (F19, F23, A34, Y36) only intra-residue or sequential NOEs are visible (except for Y36, which has NOEs to residues of the first hydrophobic core). Thus, despite transient formation of the α1-helix, there is no or only transient interactions between α1 and α2. (E) $^{15}$N spin relaxation data of ClpL NTD. Top: Longitudinal relaxation rates, middle: transverse relaxation rates, bottom: Heteronuclear NOEs. All three measurements show that the N-terminal region up to residue R35 is highly flexible, whereas the region of the hydrophobic core between α2 and the β-sheet higher heteronuclear NOE values and longer transverse relaxation rates indicating a higher degree of order. The altogether low heteronuclear NOE values, rarely above 0.6 indicate that the secondary structure elements (α1 vs. α2/β-sheet) are mobile with respect to each other in the ps-ns time scale. (F) Sequence alignment of ClpL NTDs. Similar and identical residues are highlighted in light and dark blue. The positions of secondary structural elements and of patches (A-E) composed of Y, F, N and Q residues are indicated.

**Figure 4 – figure supplement 2**

Mutant analysis of ClpL NTD. (A/D) ATPase activities of indicated ClpL mutants determined at 0.125 and 1 μM protein concentration. The ATPase activities of ClpL WT were set to 1. (B/C) Luciferase and MDH disaggregation activities (% refolded enzyme/min) of ClpL WT and indicated mutants were determined. The disaggregation activity of ClpL WT was set to 1. (E/F) Luciferase and MDH disaggregation activities (% refolded enzyme/min) of ClpL WT, $L_N$-ClpB* and $L_N$-ClpB* mutants were determined. The disaggregation activity of ClpL WT was set to 1. (G) ATPase activities of ClpL WT, $L_N$-ClpB* and $L_N$-ClpB* mutants were determined at 0.125 and 1 μM protein concentration. The disaggregation activity of ClpL WT was set to 1. (H) ClpB*, $L_N$-ClpB* and indicated mutants were incubated with aggregated MDH in presence of ATPγS. The extent of aggregate binding (% of total protein) was determined by co-sedimentation upon centrifugation. Shown are data points and mean ± SD (A-H), n ≥ 3. Statistical Analysis: One-way ANOVA, Welch's Test for post-hoc multiple comparisons. Significance levels: *p < 0.05; **p < 0.01; ***p < 0.001. n.s.: not significant.

**Figure 5 – figure supplement 1**

Multiple ClpL NTDs are required for disaggregation activity. (A) Relative Luciferase disaggregation activities (% refolded Luciferase/min) of indicated $L_N$-ClpB* disaggregase mixtures were determined (DWB: ATPase deficient $L_N$-ClpB*-E279A/E678A, ΔN: ΔN - ClpB*). Protein concentrations (µM ) are indicated. The disaggregation activity of 0.3 µM $L_N$-ClpB* was set to 1. (B) Relative Luciferase disaggregation activities (% refolded Luciferase/min) of $L_N$-ClpB* were determined in absence and presence of ΔN-ClpB* excess as indicated. The disaggregation activity of $L_N$-ClpB* was set to 1. (C) Relative Luciferase disaggregation activities (% refolded Luciferase/min (= rate)) were determined for mixtures of $L_N$-ClpB* and ΔN-ClpB* and were set as 100% for non-mixed $L_N$-ClpB* (0.3 µM). Mixing ratios are indicated as final concentration of $L_N$-ClpB* in 0.3 µM mixed hexamers. As control disaggregation activities of lower $L_N$-ClpB* concentrations were determined (+ buffer). (D) Relative Luciferase disaggregation activities (% refolded Luciferase after 120 min (= yield)) of mixed $L_N$-ClpB*/ΔN-ClpB* hexamers were determined and compared with curves calculated from a model (black to grey), which assumes that a mixed hexamer only displays disaggregation activity if it contains the number of NTDs indicated. Mixing ratios are indicated as number of ΔN-ClpB* in a hexamer. (E) Relative Luciferase disaggregation activities (% refolded Luciferase after 120 min (= yield)) were determined for mixtures of $L_N$-ClpB* and ΔN-ClpB* and were set as 100% for non-mixed $L_N$-ClpB* (0.3 µM). Mixing ratios are indicated as final concentration of $L_N$-ClpB* in 0.3 µM mixed hexamers. As control disaggregation activities of lower $L_N$-ClpB* concentrations were determined (+ buffer). Shown are data points and mean ± SD (A/B) or mean curves and datapoints (C/D/E), n ≥ 3. Statistical Analysis: One-way ANOVA, Welch's Test for post-hoc multiple comparisons. Significance levels: *p < 0.05; **p < 0.01; ***p < 0.001.

**Figure 6 – figure supplement 1**

ClpL rings interact in a M-domain dependent manner. (A) Oligomeric states of ClpL WT and indicated mutants were determined in presence of ATP by size exclusion chromatography. ATPase-deficient ClpB-E279A/E678A (DWB) served as reference for hexameric rings. Elution fractions were analyzed by SDS-PAGE. Positions of peak fractions of a protein standard are indicated. (B) Model of *Lm* ClpL highlighting the residues E352 and F354, which are crucial for M-domain mediated ring interactions. (C) Negative stain EM of indicated ClpL mutants. The insets show a magnification of a representative grid area. The scale bar is 100 nm. (D) Melting temperatures ($T_M$) of ClpL WT and indicated deletion mutants were determined in presence of ATPγS by nanoDSF or SYPRO®Orange binding. (E) Populations of diverse ClpL assembly states were determined based on 2D class averages for indicated ClpL mutants. (F) ATPase activities of indicated ClpL M-domain mutants were determined at 0.125 and 1 µM protein concentration. The ATPase activities of ClpL WT were set to 1. Standard deviations are based on at least three independent experiments (D/F). Shown data points and mean ± SD (D/F), n ≥ 3. Statistical Analysis: One-way ANOVA, Welch's Test for post-hoc multiple comparisons. Significance levels: *p < 0.05; **p < 0.01; ***p < 0.001. n.s.: not significant.

**Figure 6 – figure supplement 2**

Stabilizing ClpL ring dimers by disulfide crosslinking. (A) Relative ATPase activities of indicated ClpL WT and T355C were determined at 0.125 and 1 μM protein concentration. The ATPase activities of ClpL WT were set to 1. (B) Relative Luciferase disaggregation activities (% refolded enzyme/min) of ClpL WT and T355C were determined. The disaggregation activity of ClpL WT was set to 1. (C) Negative stain EM of ClpL T355C in oxidized and reduced states in presence of ATP. The insets show a magnification of a representative grid area. The scale bar is 100 nm. (D) YFP unfolding rates (% YFP fluorescence/min) during disaggregation of aggregated Luciferase-YFP by reduced ClpL WT (assay control) and oxidized ClpL WT (treatment control) or ClpL T355C were determined in absence of DTT (-DTT). Oxidized variants (WT and T355C) were additionally preincubated with 10 mM DTT for 30 min and tested for YFP unfolding activity in presence of DTT (+DTT). The factor of increase in YFP unfolding activity upon reduction (+DTT) is indicated (right). Shown data points and mean ± SD (A/B/D), n ≥ 3. Standard deviations for the activity gain factors (D) have been propagated from disaggregation activity standard deviations. Statistical Analysis: One-way ANOVA, Welch's Test for post-hoc multiple comparisons. Significance levels: *p < 0.05; **p < 0.01; ***p < 0.001. n.s.: not significant.

**Figure 6 – figure supplement 3**

Production of ClpL MD mutants cause increased toxicity in E. coli. (A) Production levels of ClpL (WT and indicated mutants) in *E. coli ΔclpB* cells. *E. coli ΔclpB* cells harboring plasmids for IPTG-controlled expression of *clpL* (WT and indicated mutants) were spotted on LB plates including the indicated IPTG concentrations (μM) and incubated at 30°C for 24 h. Total cell extracts were prepared from colonies and ClpL levels were determined by SDS-PAGE followed by Coomassie-staining. p: empty vector control. (B) *E. coli ΔclpB* cells harboring plasmids for IPTG-controlled expression of *clpL* (WT and indicated mutants) were spotted on LB plates including the indicated IPTG concentrations (μM) and incubated at 30°C and 42°C for 24 h.
